# Self-assembling viral histones unravel early nucleosome evolution

**DOI:** 10.1101/2023.09.20.558576

**Authors:** Nicholas A. T. Irwin, Thomas A. Richards

## Abstract

Nucleosomes are a core-component of eukaryotic nuclei, forming the structural basis of chromatin and co-ordinating processes from gene expression to chromosome segregation. Composed of a DNA-protein complex consisting of the four individual histones, H2A, H2B, H3, and H4, the nucleosome and its associated functions were key innovations during eukaryotic evolution^1,2^. However, functional constraints and the extinction of stem-eukaryotes have concealed how these dynamic systems evolved from simpler histone homologues in Archaea^3–5^. Viral histones have also previously been identified and are thought to reflect an ancestral state as they often comprise multiple histone paralogues arranged within a single protein, termed histone repeats^6–11^. Here, using viruses as an alternative source of variation, we expand the known diversity of histones and develop an empirical hypothesis for the origin of the nucleosome. Our analysis identified hundreds of histones with variable domain repeat configurations including histone singlets, doublets, triplets, and quadruplets, the latter comprising the four core histones arranged in series. Viral histone repeats consistently branch between Archaea and eukaryotes in phylogenetic trees and display intermediate functions, self-assembling into eukaryotic-like nucleosomes that stack into archaeal-like oligomers capable of impacting genomic activity and condensing DNA. The linkers conjoining the histone repeats also facilitate nucleosome formation and can promote the assembly of eukaryotic nucleosomes in the bacterium, *Escherichia coli*. Combining these data, we hypothesize that viral histone repeats represent molecular relics acquired by viruses from stem-eukaryotes during eukaryogenesis and suggest that nucleosome evolution may have proceeded through histone repeat intermediates.

---

Understanding the evolution of core eukaryotic traits, such as the nucleosome, is essential for interpreting the origin and diversification of eukaryotes^1^. However, ancestral state reconstruction is complicated by taxonomic limitations, especially the lack of extant intermediates between Archaea and eukaryotes^12^. Large double stranded DNA viruses, such as the Nucleocytoviricota viruses (NCVs), are a promising source of additional diversity as they can encode hundreds of co-opted genes with both recent and ancient cellular ancestries^13–16^. Indeed, histone proteins have been characterized from viral genomes^7,8,11^. These histones are related to each of the core histone paralogue families and exist as either individual or repeated histone-fold domains. For example, the histone doublets of marseilleviruses include H2B-H2A and H4-H3 repeats capable of forming eukaryotic-like nucleosomes essential for genome packaging^9,10,17^. Moreover, phylogenetic analyses have placed these histones between archaeal and eukaryotic homologues, prompting competing hypotheses about their origins^6,7^. In particular, whether these repeats formed by post-hoc fusion between recently transferred eukaryotic histones, represent the viral progenitors of eukaryotic histones, or were acquired following gene transfer from primordial eukaryotes, remains unclear. This ambiguity is due to under sampling of viral histone diversity, lack of phylogenetic resolution, and limited functional data.

## The diversity and phylogeny of viral histones

To differentiate between these hypotheses and to understand the relationship between viral histone repeats and nucleosome evolution, we sought to expand known viral histone diversity by surveying the proteomes of diverse NCVs, viruses known to encode histone proteins^6^. Using profile hidden Markov models representing each of the core eukaryotic histone families and archaeal histones, we searched predicted proteins derived from both NCV genomes (*n* = 205) and assembled NCV metagenomes (*n* = 2,074)^13^. This approach identified 258 complete histone genes from 168 viruses. These histones were significantly larger relative to cellular homologues, an effect explained by the presence of histone repeats such as doublets (*n* = 90), triplets (*n* = 32), and quadruplets (*n* = 13), as noted previously^6–8^ (Fig. 1a, b). These histone repeats exhibited variable but constrained domain orders, with H2A/H2B and H3/H4 nearly always in series (Fig. 1b). Similar histone repeats were also detected in ocean metatranscriptomes^18^ demonstrating that these are not artefactual gene predictions or immediately processed post-transcriptionally (Extended Data Figure 1a). Likewise, consistent post-translational cleavage sites between histone domains were absent in histone quadruplets (Extended Data Figure 1b). These data indicate that these histones are encoded and expressed as repeats.

**Figure 1.**
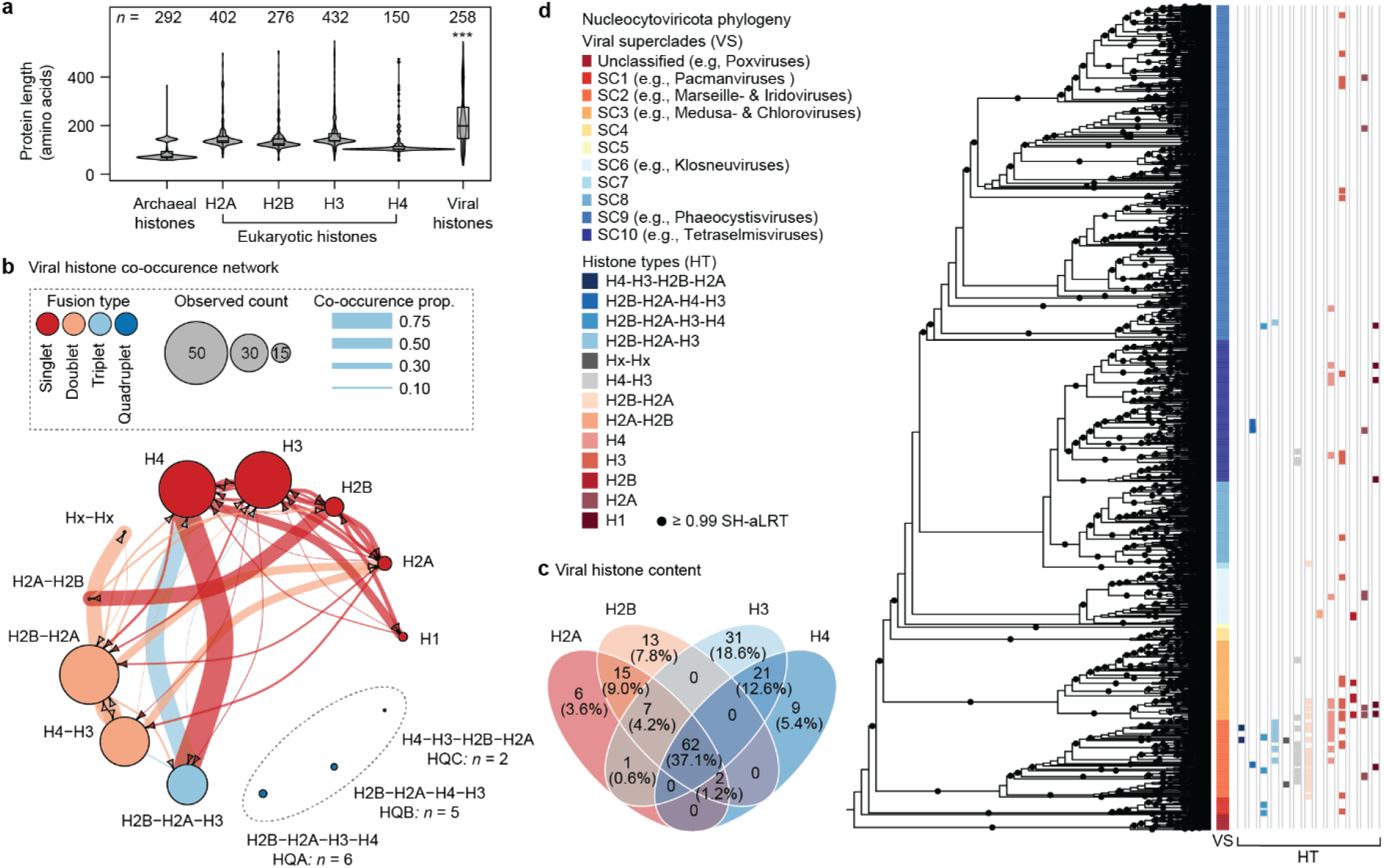
Viral histones are present and possess variable repeat domain architectures in diverse Nucleocytoviricota viruses. **a**, Amino acid length distributions of cellular and viral histones with sample sizes denoted. ***; contrast ANOVA, cellular versus viral histones, *p* = 8.47x10^-6^, F = 19.94, df = 1. **b**, A network showing the composition, frequency, and co-occurrence proportions of each viral histone. **c**, A Venn-diagram displaying histone complements in individual Nucleocytoviricota genomes. **d**, Maximum-likelihood phylogeny of the Nucleocytoviricota based on five concatenated core viral genes (major capsid protein, B family DNA polymerase, packaging ATPase, a primase-helicase, and a transcription factor) with the presence of different histone types (HT) and superclade taxonomy denoted. The phylogeny was re-run based on the alignment of Schulz et al. using the LG+F+R10 substitution model in IQ-Tree^13^.

Protein distributions across viral taxa also provided insights into histone function. Assessing histone presence over the NCV phylogeny revealed that histone repeats were common in the early-branching superclades including marseilleviruses, iridoviruses, and medusaviruses, whereas deeper-branching viral superclades mainly encoded histone singlets (Fig. 1c). Many viruses were also observed to encode multiple histone types which was reaffirmed by co-occurrence analysis (Fig. 1b, c). Regardless of genome completeness limitations, many histone-encoding viruses contained a complete histone complement, achieved through combinations of histone types such as an H2B-H2A-H3 triplet and separate H4 singlet (Fig. 1b, d). Histone quadruplets were detected in three configurations (termed HQA, HQB, and HQC) each capable of providing a complete complement but never co-occurred with other histone homologues (Fig. 1b). There were also constraints on histone composition with viruses rarely encoding an H3 or H4 in isolation with an H2A or H2B (Fig. 1d). Given these patterns we also investigated whether additional chromatin-related proteins were present in NCV genomes (Extended Data Fig. 2a). Although some putative chromatin-associated proteins were detected, there was no connection between their abundance and histone content (Extended Data Fig. 2b). This implies that although histone composition is important and constrained, viral histones likely either function autonomously or are supplemented by host-encoded factors.

Given the abundance of viral histones, we next sought to evaluate their evolutionary history. Individual histone domains extracted from the repeats as well as eukaryotic and archaeal homologues were analysed phylogenetically. The combination of improved substitution models and histone sampling resulted in a relatively well-resolved phylogeny for the entire histone superfamily which placed histones H2A and H3 as well as H2B and H4 as sisters (Fig. 2a). This fits with functional characteristics, like the tendency for H2A and H3 to diverge into histone variants, and the suggestion that H2A/H2B arose by duplication of H3/H4 precursors^1,4^. Yet although this topology was robust to model selection, the sisterhood of H2A and H3 was influenced by amino acid composition, the phylogeny was destabilized by amino acid recoding, and topology tests failed to reject alternative histone relationships (Extended Data Fig. 3, Supplementary Table S1). Unlike the overall topology, the branching of viral histones was phylogenetically robust and consistent with previous work^6,7^ (Fig. 2a, Extended Data Fig. 3). Viral histones could be assigned to specific histone families, but repeat domains almost exclusively branched outside the eukaryotic clade, unlike those which branched within eukaryotes which were almost exclusively singlets (Fig. 2a, b). Accordingly, histone repeats seem to have an ancient ancestry whereas many viral histone singlets exemplify relatively recent horizontal gene transfers from eukaryotes, as hypothesized previously^6,15^.

**Figure 2.**
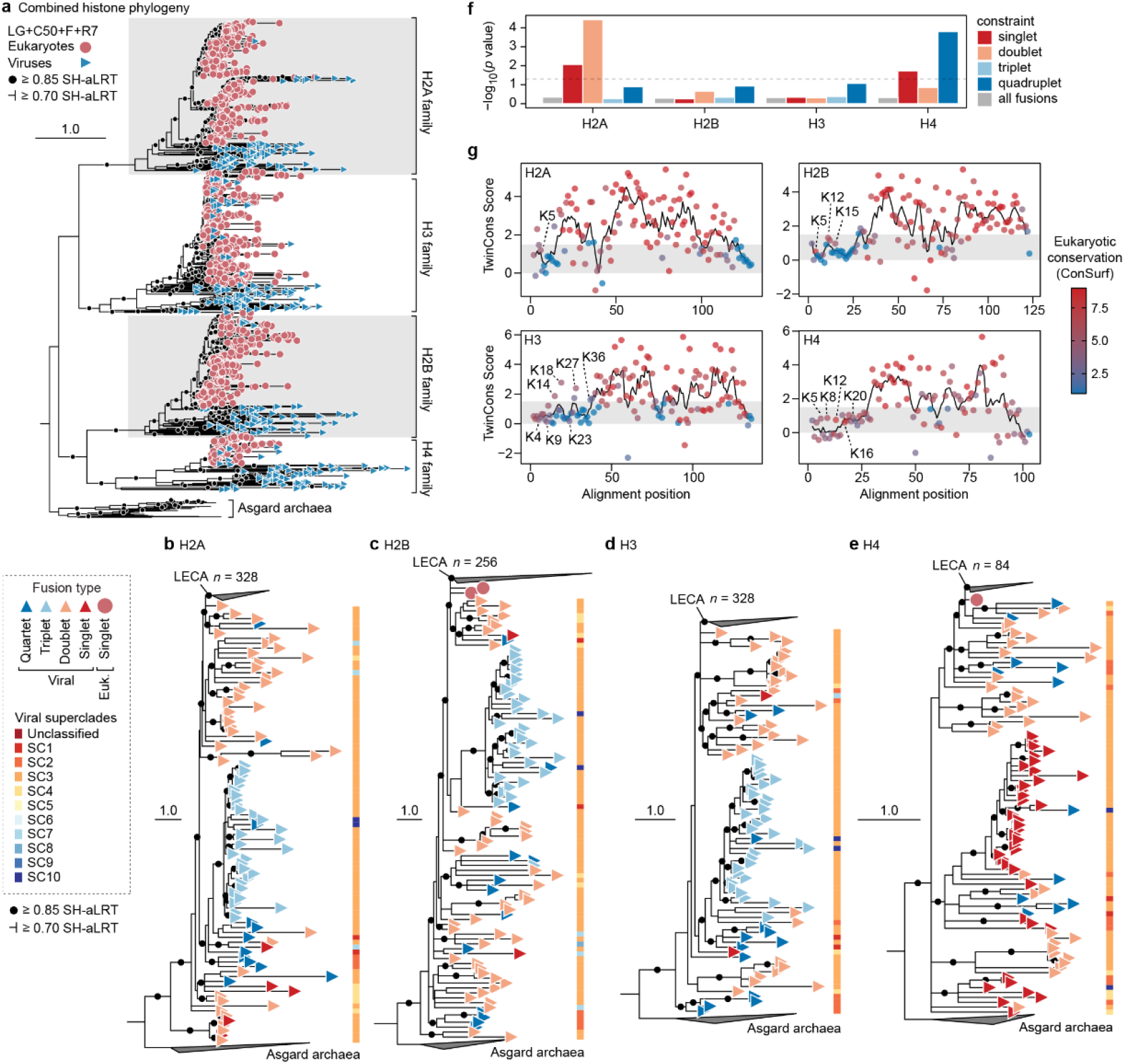
The phylogeny of viral histone repeats reveals their dynamic evolution and intermediate branching position between eukaryotes and Archaea. **a**, Maximum-likelihood (ML) phylogeny generated from an alignment of histone domains derived from eukaryotes, viruses, and Asgard archaea. Statistical support was generated using Shimodaira-Hasegawa approximate likelihood ratio tests (SH-aLRT, *n* = 1,000). **b-e**, ML phylogenies for each of the core histone families including H2A (**b**, LG+R9), H2B (**c**, LG+R7), H3 (**d**, LG+R7), and H4 (**e**, LG+R5). Eukaryotic clades (including viral histones branching within eukaryotes) have been collapsed and the node corresponding to the last eukaryotic common ancestor (LECA) is labelled. Each tree was rooted with the same histone homologues from Asgard archaea (*n* = 10). For all phylogenies, nodes with support less than 0.70 SH-aLRT were collapsed. Scale bars represent the average number of substitutions per site. Full phylogenies can be viewed at https://itol.embl.de/shared/31JevmUKUPjD4. **f**, *p* values resulting from Approximately Unbiased (AU) topology tests comparing best trees (**b-e**) to trees where the monophyly of different repeat types have been constrained. The dashed line denotes *p* = 0.05. Exact *p* values and likelihood scores are available in Table S1. **g**, TwinCons analysis comparing conservation in histone domains between eukaryotes and viral fusions. Residues are coloured based on eukaryotic histone site conservation determined using ConSurf. Dotted and dashed lines denote TwinCons values of zero and 1.5, respectively. Modifiable lysine residues have been labelled for reference.

A closer look at individual histone families revealed the relatedness and dynamic evolution of viral histone repeats (Fig. 2b-e). Histone phylogenies are difficult to fully resolve due to the time scales they cover, and the limited information provided by their short, albeit highly conserved sequences. Nonetheless, these phylogenies highlighted the nesting of eukaryotic sequences within viral homologues and illustrate histone repeat dynamics. For example, histone quadruplets often branched closely with doublets, indicating repeated fission or fusion. Similarly, the relatedness between histone quadruplets and triplets, suggest that triplets evolved following the excision of H4 from a quadruplet, although the presence of an intact quadruplet within the triplet clade could imply a reversible process. These observations were further supported by topology testing as H2A and H4 phylogenies rejected the monophyly of histone singlets, doublets, and quadruplets, but the monophyly of all histone repeats could not be rejected and these trees often placed the repeats within the eukaryotic clade (Fig. 2f, Table S1). Therefore, although these data show repeated fission and fusion during viral histone evolution, the exact order of events may be impossible to reconstruct.

To corroborate the phylogenetic data, we examined whether the evolutionary distinctiveness of viral histone repeats coincides with structural differences. To do this we compared amino acid conservation between the histone domains in eukaryotes and the orthologous domains from the viral repeats (Fig. 2g). Using TwinCons^19^, we compared amino acid conservation between the two groups. In brief, values above 1.5 approximately represent conserved sites, values below zero indicate divergent conservation, and values in between denote variability in one or both groups. To compliment this analysis, we also analysed the conservation of eukaryotic histone residues using ConSurf^20^. This analysis revealed that the histone-fold domain is conserved in both eukaryotes and viral repeats, unlike the N- and C-termini which were divergent. This included the divergence of many N-terminal sites which are conserved in eukaryotes where they are post-translationally modified and influence genomic regulation, although some N-termini contained modifiable lysines^21^. To understand the consequences of these variations, we also predicted the quaternary structures of histone quadruplets, as their lack of co-occurrence with other histones implies functional autonomy (Fig. 1b). Multimeric AlphaFold^22^ predictions indicated that histone quadruplets could assemble into dimers, in turn forming a pseudo-octameric structure with structural similarity to eukaryotic nucleosomes (Extended data Fig. 4a-c). HQA had better defined structures based on pLDDT (predicted local distance difference tests) compared to HQB and HQC (Extended data Fig. 4c-d). Regardless, histone quadruplets were predicted to form nucleosome-like structures the formation of which appeared to be facilitated by disordered amino acid linkers conjoining the histone domains (Extended data Fig. 4e) which varied in length (median lengths and standard deviations: HQA, 12 ± 5, 56 ± 6, 16 ± 8, *n* = 6; HQB, 13 ± 1, 47 ± 8, 53 ± 3, *n* = 5; HQC, 38 ± 7, 23.5 ± 1, 13.5 ± 1, *n* = 2). Some of the quadruplets also featured long disordered N- and C-terminal extensions but these were inconsistent and unconserved (Extended Data Fig. 1). The structural modelling suggest that, despite their repetitive forms and distinct evolutionary history, histone quadruplets, like histone doublets^9,10^, have the capacity to form nucleosomes.

## Viral histone quadruplets self-assemble into nucleosome-like structures

Based on the structural predictions, we tested whether viral histone quadruplets could form nucleosomes by selecting multiple representatives from each repeat configuration, alongside two archaeal histones paralogues from *Methanothermus fervidus*, which we expressed in *E. coli*, a bacterium natively lacking histones. GFP was also included as an overexpression control. Immunoblotting revealed the successful production of each protein at comparable levels, except for the archaeal histones (Fig. 3a). However, differences in molecular weight and composition make comparisons of blotting results unreliable. Nonetheless, HQA and HQC histones were largely expressed and maintained as individual proteins, whereas HQB consistently fragmented, despite attempts to avoid degradative conditions (Fig. 3a). Following expression, we investigated whether histone quadruplets could form nucleosomal structures using micrococcal nuclease (MNase), an enzyme which preferentially degrades non-nucleosomal DNA^23^. As expected, *E. coli* containing empty vectors or expressing GFP exhibited no DNA protection following MNase digestion (Fig. 3b, Extended Data Fig. 5a). In contrast, and consistent with previous results^24^, archaeal histones produced highly regular fragments with a primary size of around 70 bp, likely generated by tetramers, which increased successively by 30 bp, probably following dimer stacking (Fig. 3b, Extended Data Fig. 5a,b)^24,25^. Like archaeal histones, HQA and HQC quadruplets produced protected fragments but with an average primary size of 148 bp (141-156 bp), matching the DNA in a eukaryotic nucleosome^26^. HQB histones provided variable protection, perhaps resulting from their partial fragmentation. HQA and HQC quadruplets also generated higher molecular weight DNA fragments which increased on average by 90% (84%-97%) of the primary fragment size (Fig. 3b, Extended data Fig. 5a,b). This increase differs from the expected digestion profile of eukaryotic chromatin, where higher order fragments are produced from multiple nucleosomes and linker DNA (often between 20 and 80 bp). This difference suggests that rather than forming typical nucleosomal arrays, histone quadruplets may oligomerize without linker DNA, similar to how archaeal histones function^25^. To understand the structure of these putative oligomers, we covalently crosslinked histones *in vivo* and assessed their molecular weights using immunoblotting (Fig. 3c). As predicted, archaeal histone HmfA and histone quadruplets HQA2, HQB2, and HQC2 each displayed higher order bands following crosslinking, with molecular weights approximately double or triple that of the monomers, consistent with oligomerization rather than non-specific binding. This implies that histone quadruplets form dimers, in agreement with the structural predictions, and potentially higher order structures (e.g., HQA2) which would account for the digestion profiles. Although additional structural work will be required to fully understand the nature of these proteins, these data indicate that histone quadruplets can be expressed as complete repeats, assemble without native chromatin machinery, dimerize, and bind DNA forming eukaryotic-like nucleosomes that stack into archaeal-like oligomers, possibly through dimer addition (Fig. 3d).

**Figure 3.**
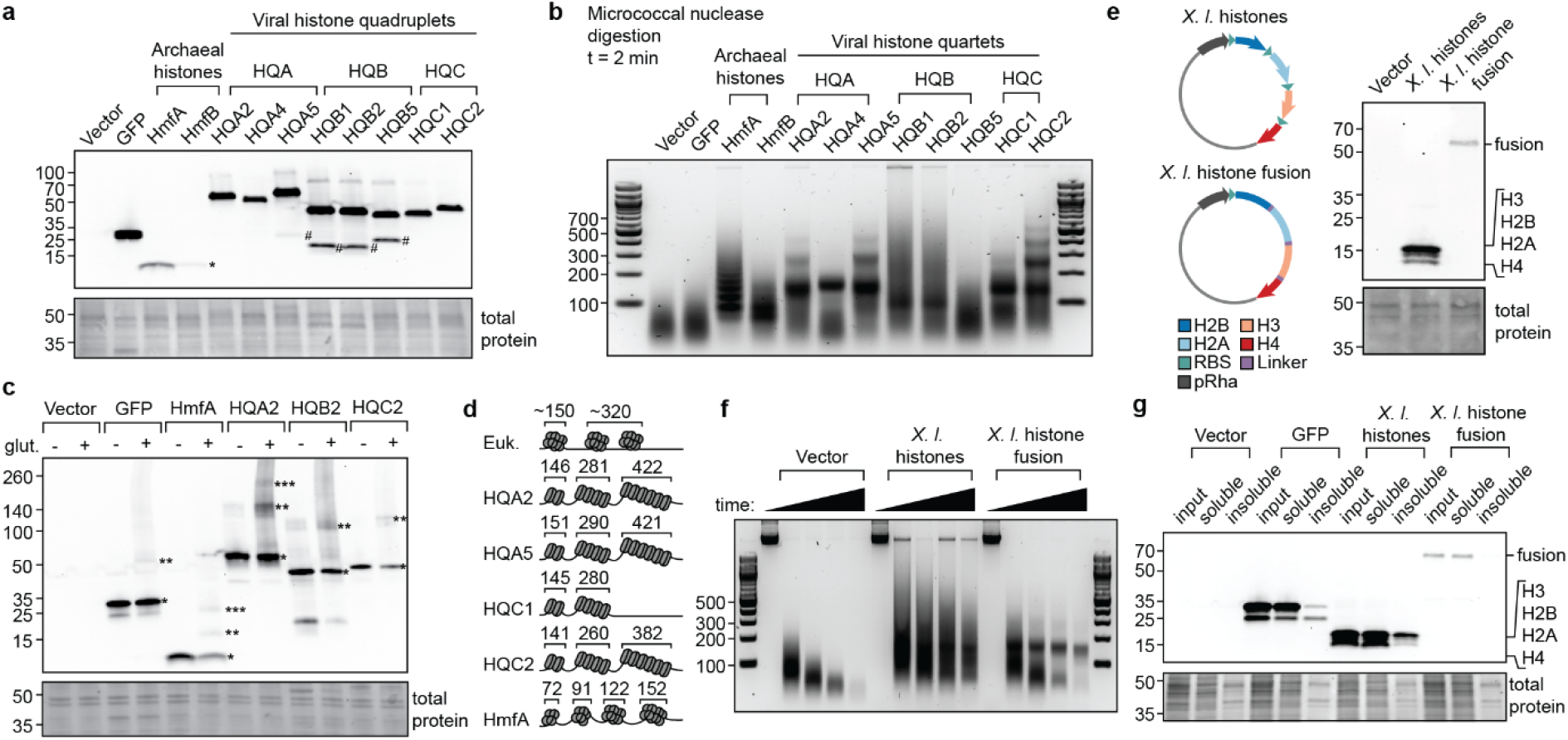
Viral histone quadruplets self-assemble into nucleosomal structures. **a**, anti-6xHis immunoblot on whole cell extracts from induced *E. coli*. Degradation products (#) and archaeal HmfB (*) are denoted. **b**, Gel electrophoresis of genomic DNA from *E. coli* after two minutes of micrococcal nuclease (MNase) digestion. **c**, anti-6xHis immunoblot of whole cell extracts with and without glutaraldehyde-crosslinking. Putative oligomers are highlighted with asterisks. Note that some higher molecular weight products are still recovered without cross-linking (e.g., HQA2 and HQB2) despite denaturing conditions. **d**, Hypothetical model for nucleosome formation with eukaryotic, archaeal, and viral histones. **e**, Diagram of *Xenopus laevis* (*X. l*.) histone constructs and anti-6xHis immunoblot of whole cell extracts from *E. coli* expressing each construct. RBS, ribosome binding site. **f**, Gel electrophoresis of genomic DNA from *E. coli* after 1, 2, 5, and 15 minutes of MNase digestion while expressing *X. laevis* histone vectors. **g**, Anti-6xHis immunoblot showing input, soluble, and insoluble protein fractions from *E. coli* expressing GFP or *X. laevis* histones induced by 1 mM rhamnose. Each experiment was repeated at least twice independently with equivalent results.

Nucleosome assembly in eukaryotes typically requires chaperones (e.g., NAP1, ASF1, CAF1^27^), yet viral histone quadruplets formed on the *E. coli* genome without native proteins. To understand how histone quadruplets assemble, we tested the role of the amino acid linkers in nucleosome formation by generating two constructs containing the four core histones from the frog, *Xenopus laevis* (Fig. 3e). These histones were either expressed polycistronically in stoichiometric amounts using repeated ribosome binding sites^28^ or were conjoined using HQA1 linkers. Both constructs were expressed in *E. coli* and protein production was confirmed by immunoblot (Fig. 3e). Micrococcal nuclease digestions revealed that both separated and fused *X. laevis* histones protected DNA. However, when separated, the DNA footprint was varied and smeared, similar to HQB quadruplets (Fig.3f). In contrast, fused *X. laevis* histones generated a defined singular band at 152 bp in accordance with canonical nucleosomes (Fig. 3f, Extended data fig. 5c). This fragment was also present but less apparent when histones were separated. These experiments indicate that eukaryotic histones can assemble into nucleosomes without native chaperones, but fusion appears to facilitate assembly, perhaps by improving folding, stoichiometry, and binding consistency. Protein misfolding in *E. coli* typically results in aggregation and incorporation into insoluble inclusion bodies^29^. Therefore, we inspected the solubility of these histones when expressed at varying levels by adjusting rhamnose inducer concentrations (Fig. 3g, Extended data fig. 5d,e). Although separated histones, alongside GFP, were consistently detected in the insoluble fraction, the fused histone was nearly absent. Moreover, GFP insolubility was dependent on expression level unlike the separated histones which were consistently insoluble (Extended data fig. 5d,e). This suggests that histone insolubility is a result of ineffective folding or assembly rather than overexpression, an effect mitigated by fusion.

Given the ability of histone quadruplets to assemble on the *E. coli* genome and the known regulatory capacity of nucleosomes^2^, we assessed the phenotypic impact of histone expression on genomic function in *E. coli*. Under standard conditions, histone expression had a small or undetectable effect on maximum growth (A) and growth rate (μ) but increased lag-phase (λ), suggesting a disruption to the transition out of stationary phase, rather than the cell cycle (Fig. 4a, Extended Data Figure 6). Challenging histone expressing strains with antibiotics targeting genomic activities further altered phenotypes. Histone expression resulted in high sensitivity to novobiocin, a DNA gyrase inhibitor that causes excessive negative supercoiling, as well as zeocin, which induces double stranded DNA breaks (Fig. 4b,c). Drug-challenged strains exhibited impaired growth but had normal lag-phases relative to uninduced cultures. These effects were less pronounced in the presence of rifampicin, a transcription inhibitor, where growth dynamics mirrored the control (Fig. 4d). However, low concentrations of rifampicin were used to mediate transcriptional defects. Regardless, these synthetic lethal phenotypes indicate that histone quadruplets, like archaeal and *X. laevis* histones, impact genomic function likely by relieving negative supercoiling, a canonical nucleosomal function^30^, and impairing DNA repair, perhaps by reducing DNA accessibility.

**Figure 4.**
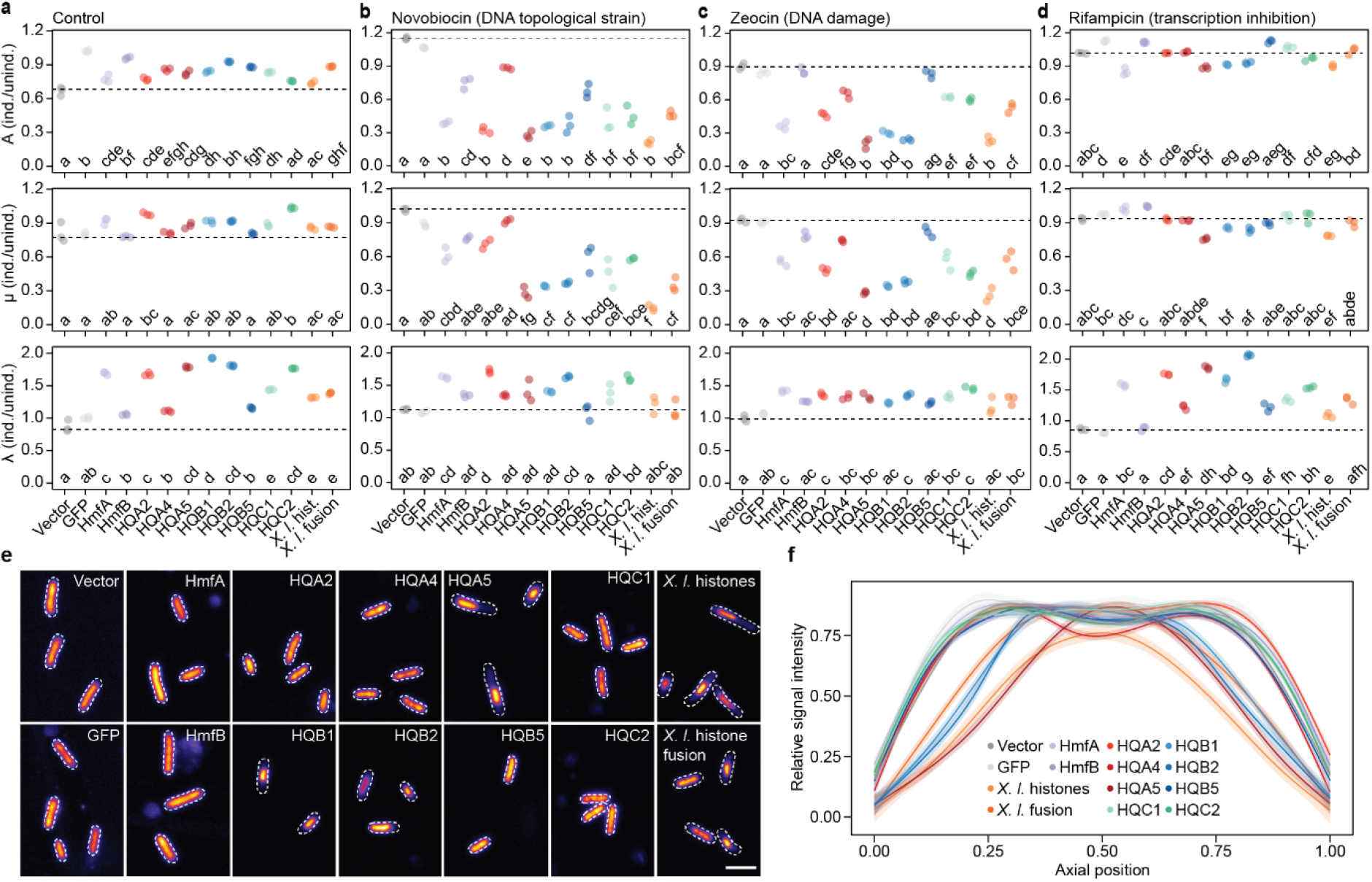
Histone quadruplets impact genomic function and nucleoid morphology in *Escherichia coli*. Growth characteristics inferred from OD_600_ measurements including maximum growth (A), maximum growth rate (μ), and lag-phase (λ) in induced compared to uninduced strains. Growth parameters were inferred for three biological replicates under standard conditions (**a**) or with the addition of 150 μM novobiocin (**b**), 1 μM zeocin (**c**), or 1 μM rifampicin (**d**). Conditions were compared using pairwise Tukey HSD (Honestly Significant Difference) tests and significance groups are denoted using a compact letter display (*p* < 0.01 after Bonferroni multiple test correction). See Extended Data Figure 6 for full growth curves. **e**, Representative confocal fluorescent micrographs of *E. coli* cells after staining with Hoescht. Images have been coloured using the Fire LUT (Lookup table) in ImageJ to show fluorescent intensity and cell outlines are marked with dashed lines, based on brightfield microscopy. Scale bar: 2 μm. **f**, Hoescht staining signal intensity measured along the long axis of *E. coli* cells, normalized to the maximum intensity value, and averaged across individuals (*n* = 23, per strain). Shaded regions represent standard error. Only cells of consistent length (2-4 μm) were compared (ANOVA, *p* = 0.103, F = 1.535, df = 13). Each experiment was repeated at least twice independently with equivalent results.

Histone-dependent phenotypic effects also manifested as altered nucleoid morphology. In the presence of histone quadruplets HQA5, HQB1, and HQB2, as well as separated and fused *X. laevis* histones, *E. coli* nucleoids were condensed (Fig. 4e,f). This effect was not observed for other viral quadruplets or archaeal histones. Growth phenotypes, MNase digestion profiles, and nucleoid condensation roughly correlated (i.e., strains with more DNA protection exhibited stronger growth defects and DNA compaction), although some quadruplets with well-defined digestion profiles (e.g., HQA2, HQC1, and HQC2) did not alter nucleoid structure. These data indicate that amino acid sequence influences histone repeat function more than domain order and reveals another intermediate feature as some histone quadruplets induce eukaryotic-like DNA condensation whereas others do not, like archaeal histones^24^. However, additional experimentation with other eukaryotic and archaeal histones could reveal more functional variability.

## The evolution of viral histone repeats and the origin of the nucleosome

Here we expand the known diversity of viral histones and further resolve the wider histone phylogeny corroborating previous analyses hinting at the ancient evolutionary history of histone repeats^6,7^. We further demonstrate that histone quadruplets can self-assemble on a naïve genome where they form eukaryotic-like nucleosomes, stacking into archaeal-like oligomers that affect genomic function and nucleoid morphology. The successful formation of viral and eukaryotic nucleosomes in *E. coli* provides new experimental possibilities for synthetic chromatin biology.

The evolutionary and functional results reported here also allow us to explore the origins of viral histone repeats and the nucleosome more generally. Interpreting the evolution of these proteins requires evaluating three alternative origin hypotheses. Firstly, the formation of histone repeats could have followed the fusion of eukaryotic histones acquired by horizontal gene transfer to viruses after the last eukaryotic common ancestor (LECA). This proposition is inconsistent with histone phylogenies, despite improved sampling and phylogenetic resolution, as histone repeats consistently branch intermediately between archaea and eukaryotes (Fig. 2a-e). Conserved N-terminal residues are also largely absent from the repeats, necessitating significant divergence across multiple domains after acquisition. Furthermore, eukaryotic histones were rarely observed in repeat architectures implying that histone fusion is uncommon in extant eukaryotic lineages. Secondly, viral histone repeats could represent the progenitors of eukaryotic histones. This fits phylogenetically but the absence of additional chromatin proteins in NCV genomes indicates two unlikely scenarios where eukaryotes either evolved regulatory machinery after histone acquisition or viral chromatin proteins were transferred to a eukaryotic ancestor before being comprehensively lost in NCV lineages. Thirdly, histone repeats could have been acquired by NCV viruses from stem-eukaryotes during or just prior to the emergence of LECA (Fig. 5). This is consistent with the phylogenetic results and with the intermediate functional characteristics of the quadruplets. Likewise, sequence divergence and the lack of additional chromatin proteins could reflect an ancestral proto-eukaryotic state and host-dependency, respectively. If correct, these histones could represent molecular relics revealing snapshots of histone evolution during eukaryogenesis. These proteins have evidently changed over time and their original contexts have been lost. Nonetheless, the form and function of viral histone repeats can help dissect nucleosome evolution and if similar gene transfers have occurred for other proteins, these events could help resolve key aspects of the black box of eukaryogenesis.

**Figure 5.**
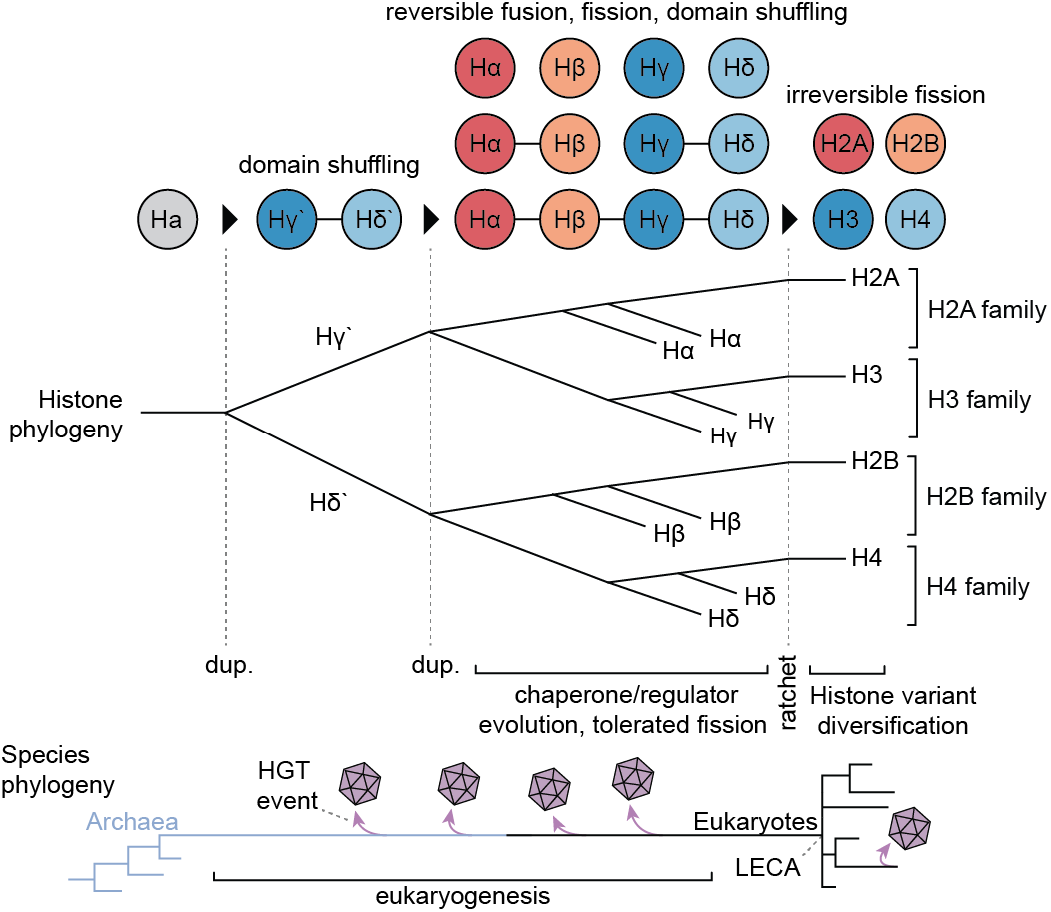
A hypothesis for the origin of the nucleosome. The top and bottom phylogenies illustrate hypothesized evolutionary histories for histones and species during eukaryogenesis, respectively. Key events in histone evolution are noted including duplications (dup.) and an evolutionary ratchet, locking eukaryotic histones in their individualized states (e.g., an evolved dependency on N-terminal tail modifications). Horizontal gene transfer (HGT) events are illustrated on the species phylogeny to demonstrate where viral histone repeats could have emerged from.

Regardless of how viral histone repeats evolved, their contributions to understanding histone function and phylogeny point to a hypothesis for the origin of the nucleosome (Fig. 5). The capacity for histone linkers to facilitate folding provides a resolution to the paradox of whether regulatory machinery or the nucleosome itself came first. We therefore hypothesize that the eukaryotic histones evolved through the repeated duplication of an archaeal-like homologue through repeat-intermediates. Based on our best histone phylogeny, the progenitor likely initially duplicated into a primordial H3 (Hγ’) and H4 (Hδ’). H3 and H4 are thought to have emerged first given their ability to form tetramers and their initializing role in nucleosome formation^4^. The infrequent co-occurrence of mixed pairs in viral genomes also suggests functional incompatibility indicating that a heterodimeric pair probably emerged first. The subsequent duplication of an Hγ’/ Hδ’ repeat would have formed the H2A/H3 and H2B/H4 families with pre-LECA representatives (Hα/Hγ, Hβ/Hδ, respectively^7^) (Fig. 5). We suggest that these histones existed as domain repeats, cooperatively assembling while chaperones and regulators evolved to facilitate this established process. Chaperone evolution would have permitted reversible fragmentation potentiating histone fusion and fission. However, histone fusions have not been observed in eukaryotes. Therefore, we argue that an evolutionary ratchet emerged prior to LECA, such as a dependency on N-terminal tails or histone variants, which was incompatible with fusion, locking eukaryotic histones into the separate proteins we observe today. Indeed, some Asgard archaeal histones contain lysine-rich N-terminal tails and post-translational modifications can regulate archaeal chromatin, indicating that tail usage has been experimented with throughout histone evolution^31,32^. Ultimately, this scenario is consistent with comparative genomic, phylogenetic, and functional data and provides a parsimonious explanation for the origin of the nucleosome, a fundamental structure which impacted the emergence of core eukaryotic traits, from linear chromosomes to complex genome regulation, and sex.

## Methods

### Histone identification and classification

To identify histone proteins in viral genomes, we developed profile hidden Markov models (HMM) for each of the five eukaryotic histone families (including H1) and archaeal histones. To this end, we assembled a taxonomically balanced set of genome-predicted proteomes from UniProt^33^ (version 2021_03) by selecting the best proteome per genus from eukaryotes (*n* = 106) and Archaea (*n* = 230) based on BUSCO (Benchmarking Universal Single Copy Orthologues) presence^34^. For opisthokonts (Metazoa and Fungi) and streptophytes, stricter criteria were used by selecting the best proteome per phyla (*n* = 24) and family (*n* = 11), respectively. The archaeal proteomes were also supplemented with additional non-reference Asgard proteomes from UniProt (*n* = 14) and the resulting proteomes from all species were separately clustered at 95% identity using CD-HIT v4.8.1^35^.

Using histone orthologues from *Homo sapiens* (H1: P07305, H2A: Q96QV6, H2B: P682807, H3: P68431, H4: P62805) and *Methanothermus fervidis* (P48781) as separate queries, we searched the proteomes using Diamond BLASTp v2.0.9 (ultra-sensitive mode, E < 10^-5^)^36,37^. The BLAST hits were extracted and aligned using the structurally informed MAFFT-DASH v7.490 multiple sequence aligner and the L-INS-i algorithm, before being trimmed to a gap-threshold of 20% using trimAl v.1.4.rev15^38,39^. Phylogenies were generated from the trimmed alignments using IQ-Tree v2.1.2 and LG (Le-Gascuel) substitution models were selected using ModelFinder^40^. The trees were inspected using FigTree v1.4 (http://tree.bio.ed.ac.uk/software/figtree/) and correct orthologues were identified and extracted based on phylogenetic topologies and sequence annotation using SWISS-PROT^41^. The identified orthologues were then re-aligned using MAFFT-DASH and used to generate HMMs with HMMER v3.1b2^42^. The HMMs were used to search the proteomes twice iteratively using HMMER (E< 10^-5^) and between each search, the hits were extracted, aligned with the previously identified homologues, phylogenetically curated, and incorporated into improved HMMs representing each of the histone families and encompassing diversity from across eukaryotes and Archaea.

Each of the histone HMMs was subsequently used to search a database of proteins predicted from diverse Nucleocytoviricota genomes and metagenomes^13^. To filter out incomplete or fragmented sequences, proteins were re-predicted from viral genomes using Prodigal v2.6.3 and annotated as complete given the presence of a start and stop codon^43^. The resulting full-length viral histone homologues evidently included fusion proteins. Therefore, to classify the histone domains found within individual proteins, the sequences were annotated with the histone HMMs using HMMER. When multiple HMMs overlapped by more than 50%, the domain with the better conditional E-value was assigned to the region and domains mapping with less than 25% HMM coverage were excluded. Annotated histone domains were then extracted and concatenated with reference sequences from each of eukaryotic and archaeal histone families before sequence alignment and phylogenetic analysis, as described above. Histone domains were annotated phylogenetically as belonging to the archaeal, H2A, H2B, H3, or H4 families. If a histone domain could not be assigned due to an intermediate branching position it was denoted Hx. The final histone types were classified based on histone domain order and composition and, to avoid prediction artifacts, only histone types observed in more than one separate genome were analysed further.

To corroborate the genome predictions, we also searched for viral histone homologues in ocean metatranscriptomes produced as part of the Tara Oceans Project^18^. To reduce database size, unigenes annotated with a histone fold domain based on Pfam (PF00125) were extracted. Transcripts were translated using TransDecoder v5.5.0 (https://github.com/TransDecoder/TransDecoder) and the predicted proteins were searched and annotated as above using the histone HMMs.

### Reconstructing chromatin processing machinery in the Nucleocytoviricota

To understand co-occurrence between histone proteins and chromatin processing machinery in the Nucleocytoviricota, we searched Nucleocytoviricota genomes and metagenomes for a suite of previously curated chromatin-associated proteins^44^. These proteins included post-translational modifiers, chaperones, readers, remodellers, and other proteins involved in complex formation from diverse eukaryotes. Previously identified orthologues were downloaded, aligned using the L-INS-i algorithm in MAFFT v7.490, and HMMs were generated from the resulting alignments. Nucleocytoviricota proteomes were then searched using these HMMs with HMMER (E < 10^-5^), and if a single protein was identified by multiple HMMs it was assigned to the HMM that produced the lower E-value. The resulting hits were visualized using a heatmap and hierarchical clustering was conducted using a correlation distance metric and the ward D2 clustering method implemented in Pvclust in R v4.2.0^45^.

### Phylogenetics and topology testing

Phylogenies were generated for individual histone families and the entire histone superfamily using IQ-Tree. Individual histone domains were extracted from repeat proteins as described above and aligned with individual eukaryotic and archaeal histones using MAFFT-DASH v7.490 and the L-INS-i algorithm. The resulting alignments were trimmed with a gap-threshold of 20% and sequences with less than 50% of sites present were removed. Fast-evolving eukaryotic taxa including parasites and amoebozoans were excluded from the analyses due to phylogenetic ambiguity (e.g., *Trichomonas, Naegleria, Dictyostelium*). For each phylogeny, substitution models were selected using ModelFinder and statistical support was generated using Shimodaira-Hasegawa approximate likelihood ratio tests (SH-aLRT, *n* = 1,000)^46^. Given the short alignments (∼100 amino acids), we used a reduced perturbation strength (-pers 0.2), an increased threshold for unsuccessful tree search iterations (-nstop 200) and ran each tree search 100 times independently to avoid local likelihood maxima, selecting the best tree from each set of replicates. Note that phylogenies calculated using empirical profile mixture models were only ran once due to computational limitations. To account for phylogenetic artifacts including long-branch attraction and compositional bias, phylogenies were re-ran using alternative substitution models, 4-state Dayhoff recoded alignments, and alignments lacking compositionally biased sequences based on composition χ^2^ tests (*p* < 0.05) performed in IQ-Tree^47^. For topology testing, topological constraints were applied and constrained trees were calculated as above and compared to the best unconstrained tree using Approximately Unbiased (AU) tests^48^. For testing the monophyly of proteins with different domain configurations, only viral sequences branching outside of the eukaryotic clade in the best tree were constrained as monophyletic. Phylogenies were visualized in IToL v6^49^.

### Sequence and structural analysis

Histone repeat sequences were analysed computationally to understand functional, structural, and evolutionary characteristics. Predicted proteolytic cleavage sites were assessed using the PeptideCutter tool in ExPasy whereas protein tertiary structure was calculated using AlphaFold v2.3.2 implemented through ColabFold^22,50,51^. For the AlphaFold predictions, structures were predicted using the AlphaFold 2-multimer-v2 model given a dimeric configuration with multiple sequence alignments generated using homologues from the MMSeqs2 database^52^. Protein structure visualization and root mean square deviation calculations (RMSD) were done using Pymol (https://github.com/schrodinger/pymol-open-source). DNA was added to the models from a *Xenopus laevis* nucleosome structure (PDB: 6ESF), following structural alignment in Pymol. To investigate eukaryotic sequence conservation, multiple sequence alignments and phylogenies for each histone family (excluding viral sequences) were analysed using ConSurf^20^. Conservation comparisons between viral and eukaryotic sequences were done using TwinCons using the LG substitution model and Voronoi clustering^19^.

### Histone expression and immunoblotting

Archaeal, viral, and eukaryotic histone genes with C-terminal 6xHis tags were expressed in *Escherichia coli* strain K12 from a rhamnose-inducible promoter (pRha) in a pD861 vector after being codon optimized and synthesized by Synbio Technologies. *Xenopus laevis* histones (P06897, P02281, P02302, P62799) were either expressed polycistronically with each open reading frame separated by a ribosome binding site or were joined with three linkers from histone quadruplet HQA1 (AEAEDVK, TELGELINKQLFNDDRKRKLATARRNRRKAEDTGDADGATTSG, QPARLNT)^28^. Plasmids were transformed into *E. coli* by heat shock and were maintained using kanamycin selection (50 μg/mL). To induce histone expression, cells were grown overnight in LB (lysogeny broth) at 37 °C before being diluted 1:100 into fresh LB and grown at 30 °C to an OD_600_ (optical density at 600 nm) of 0.6. Prior to induction, cultures were cold shocked in an ice water bath for 30 minutes while shaking at 100 rpm to improve protein solubility after induction. The inducer, L-rhamnose monohydrate, was then added to 10 mM (unless otherwise noted) and cultures were induced for 18 hours at 20 °C, shaking at 180 rpm.

Immunoblots were used to confirm protein expression and assess histone fusion properties. Total protein extracts were collected by centrifuging 150 μL of induced culture at 15,000 xg for two minutes, resuspending the cell pellet to an OD_600_ of 1.0 in SDS loading buffer (50 mM Tris pH 6.8, 2% SDS, 10% glycerol, 100 mM β-mercaptoethanol), and incubating the extracts at 37 °C for 10 minutes. Protein extracts were loaded into 4-20% Tris-glycine SDS-PAGE gels (BioRad) and ran at 120 V for 30 minutes in Tris-glycine-SDS buffer (2.5 mM Tris, 19.2 mM glycine, 0.01% SDS, pH 8.3). Polyacrylamide gels were equilibrated for 15 minutes in transfer buffer (2.5 mM Tris, 19.2 mM glycine, 20% methanol, pH 8.3) at room temperature and then transferred onto 0.2 μm pore-size nitrocellulose membranes at 30 V for 70 minutes at 4 °C. Total transferred protein was assessed using Pierce™ Reversible Protein Stain (Thermo Fischer) and membranes were blocked for two hours at room temperature with 0.2% alkali soluble casein in PBST (10 mM phosphate, 150 mM NaCl, 0.1% Tween-20, pH 7.5). Blocked membranes were incubated for one hour at room temperature in 0.2 μg/mL Anti-6x His-tag (ab9108) in PBS-Tween after conjugating the antibody to horseradish peroxidase (HRP) using an HRP-conjugation kit (ab102890). Finally, membranes were washed in PBST and chemiluminescence was visualized using Pierce ECL (enhanced chemiluminescence) substrate and imaged using an Invitrogen iBright CL1500.

### Micrococcal nuclease digestions

To assess nucleosome formation, *E. coli* cultures were induced, fixed, lysed, and digested with micrococcal nuclease, an enzyme that preferentially digests non-nucleosomal DNA, based on previous methods^24^. Induced cultures (10 mL) were prepared as described above, before being fixed in 1% formaldehyde for 15 minutes while shaking at 180 rpm. Fixation was quenched with 125 mM glycine for five minutes and the resulting cells were pelleted, washed in cold PBS, and stored at -80 °C. Thawed cell pellets were then resuspended and digested for an hour on ice with freshly made lysozyme buffer (120 mM Tris pH 8, 50 mM EDTA, 4 mg/mL lysozyme (Thermo Fischer 89833)). Protoplasts were pelleted at 15,000 xg for 3 minutes and resuspended in 500 μL of cold lysis buffer (10 mM NaCl, 10 mM Tris pH 8, 3 mM MgCl2, 0.5% NP-40, 0.15 mM spermine, 0.5 mM spermidine, Roche EDTA-free protease inhibitor) and incubated on ice for 20 minutes. Lysates were pelleted at 15,000 xg for 10 minutes and the resulting pellets were gently washed in -CA buffer (15 mM NaCl, 10 mM Tris pH 7.4, 60 mM KCl, 0.15 mM spermine, 0.5 mM spermidine, EDTA-free protease inhibitor) without resuspension. Lysates were then centrifuged at 15,000 xg for 5 minutes and resuspended in 500 μL of cold +CA buffer (15 mM NaCl, 10 mM Tris pH 7.4, 5 mM CaCl2, 60 mM KCl, 0.15 mM spermine, 0.5 mM spermidine, EDTA-free protease inhibitor). Prior to digestion, samples were equilibrated to 37 °C before micrococcal nuclease (Thermo Fisher 88216) was added to 250 U/mL. Lysates were then briefly vortexed and incubated at 37 °C with samples being taken at designated time points. To stop the reaction, 0.25 volumes of STOP solution (200 mM EDTA pH 8, 200 mM EGTA pH 8) were added after sample collection. Lastly, digested samples were treated with 400 μg/mL RNaseA (Thermo Fisher EN0531) for 30 minutes at 37 °C followed by 1% SDS and 800 μg/mL proteinase K (Thermo Fisher EO491) for 16 hours at 65 °C. DNA was purified using a Qiagen PCR purification kit and samples were analysed using 2.5% agarose TBE gels (90 mM Tris, 90 mM boric acid, 2 mM EDTA pH 8) which were electrophoresed at 150V for 30 minutes. DNA fragment sizes were quantified using an Agilent 2100 Bioanalyzer with a high sensitivity DNA kit.

### Glutaraldehyde crosslinking

Induced cultures (10 mL) were prepared, and the cells pelleted and washed with cold PBS before being frozen at -80 °C. Thawed cell pellets were resuspended and digested in 1 mL of fresh lysozyme buffer on ice for one hour to generate protoplasts. The cells were then resuspended in 500 μL of cold lysis buffer (10 mM NaCl, 10 mM HEPES pH 7.5, 3 mM MgCl2, 0.5% NP-40, 0.15 mM spermine, 0.5 mM spermidine, EDTA-free protease inhibitor) using a 22-gauge syringe and left to incubate on ice for 20 minutes. Lysates were centrifuged at 15,000 xg for 10 minutes and the supernatant was removed before the pellet was again resuspended in 500 μL cold HEPES buffer (10 mM NaCl, 10 mM HEPES pH 7.5, 30 mM KCl, 3 mM MgCl2, 0.15 mM spermine, 0.5 mM spermidine, EDTA-free protease inhibitor) using a 22-gauge syringe. Samples were incubated at 20 °C for five minutes and split into 95 μL aliquots to which either water or glutaraldehyde was added to a final concentration of 0.025%. The samples were fixed at room temperature for five minutes before being quenched with 100 mM Tris pH 7.6. Fixed samples were then treated with 400 μg/mL RNaseA for 30 minutes at 37 °C and 2.5 units of DNAseI (Thermo Fisher EN0525) for one hour at 37 °C. SDS loading buffer was added to the samples which were denatured at 95 °C for 10 minutes before being analysed by SDS-PAGE and immunoblotting, as described above, but with prolonged electrophoresis and transfer times of one hour.

### Histone solubility assays

To assess the influence of histone linkers on protein solubility, *E. coli* strains (10 mL cultures) expressing fused or separated *Xenopus* histones were induced and frozen as described. To lyse the cells, cell pellets were resuspended and incubated in lysozyme buffer for one hour on ice. Protoplasts were then pelleted at 15,000 xg for 3 minutes at 4 °C and resuspended in 500 μL of cold HEPES lysis buffer. To ensure efficient lysis, the resuspended cells were then sonicated three times using a Soniprep 150 sonicator (MSE) at 50% power (10 amplitude microns) for 10 seconds with a one-minute ice incubation between each cycle. NP-40 was then added to 0.5% and the lysates were mixed well and incubated on ice for 20 minutes. An input sample was then taken, and the remaining lysates were centrifuged at 16,000 xg for 20 minutes at 4 °C. The soluble supernatant was isolated, and the remaining insoluble pellet was washed once with PBS. The insoluble pellet was then resuspended in lysis buffer and sampled. Protein samples were mixed with SDS loading buffer, denatured at 95 °C for 10 minutes, and analysed by SDS-PAGE and immunoblot.

### Growth curves and phenotyping

To investigate the phenotypic impacts of histone expression in *E. coli*, strains were inoculated and grown overnight in 5 mL of LB with kanamycin (LB-Kan). The next day, cultures were diluted 1:100 into 5 mL of fresh LB-Kan and grown to an OD_600_ of 0.6 at 37 °C. The cultures were then cold shocked in an ice water bath for 30 minutes while shaking before rhamnose-monohydrate was added to a concentration of 10 mM. Induced cultures were then grown at 30 °C for four hours, shaking at 225 rpm, before being inoculated to an initial OD_600_ of 0.025 in the wells of a 96-well plate containing 200 μL of LB-Kan supplemented with either novobiocin (150 μg/mL), rifampicin (1 μg/mL), or zeocin (1 μg/mL). Growth was monitored at 30 °C with 400 rpm double orbital shaking using a FLUOstar Omega (BMG Labtech) plate reader with measurements conducted every 30 minutes at 600 nm. The resulting data were collected and the OD_600_ of uninoculated media was subtracted from each measurement. Sigmoidal growth curves were then fit to individual biological replicates using a non-linear least square fitting function (maximum iterations = 10,000, tolerance = 1x10^-5^) based on the formula:

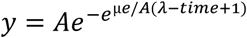

where μ, A, and λ represent the maximum growth rate defined as the maximum slope, maximum growth interpreted as the curve maximum, and the lag-phase time determined as the time of the maximum slope, respectively. All analyses were conducted in R v4.2.0.

### Nucleoid imaging

Induced *E. coli* strains were prepared as described above, fixed with 0.2% glutaraldehyde, and incubated with 15 μg/mL Hoescht 33342 for 30 minutes in the dark at room temperature. Stained cells were then applied to a slide and mixed with an equal volume of 50% CitiFluor AF2 antifade solution in PBS. Slides were sealed and imaged using a Zeiss LSM-780 inverted high-resolution laser scanning confocal microscope with a Ph3 ×100 oil objective. Exposures were kept constant during experiments, and images were collected using BLACK ZEN Software (ZEN Digital Imaging for Light Microscopy) before being analysed with ImageJ (https://imagej.net). Nucleoid density was measured in ImageJ by measuring the fluorescent signal intensity along a transect between each cell end, determined using brightfield microscopy. Signal intensity was made relative to the maximum measured value and was averaged across measured cells. Only cells which were clearly non-dividing and lying flat in the focal plane were measured. Likewise, to control for variation in cell size, only cells between 2-4 μm were included in the analysis.

## Acknowledgements

We thank Tobias Warnecke and Antoine Hocher for discussions, suggestions, and expression vectors, as well as Luis Javier Galindo Gonzaléz for assistance with fluorescent microscopy. N.A.T.I. was supported by a Junior Research Fellowship from Merton College, Oxford. T.A.R. was supported by a Royal Society University Research Fellowship (URF\R\191005) and Leverhulme Grant (RPG-2014-054).

## Author contributions

N.A.T.I. conceptualized the project, conducted the experiments, analysed the data, and wrote the manuscript. T.A.R. provided support and edited the manuscript.

## Materials and Correspondence

All data including proteomes, HMMs, alignments, phylogenies, and identified NCV chromatin proteins, as well as unedited gel and blot images, unprocessed micrographs, and expression construct sequences are available from FigShare (https://figshare.com/s/40c5ee5552097be43c6b). Expression vectors and *E. coli* strains are available upon request. Requests and correspondence can be addressed to N.A.T.I.

## Supplemental Figures

**Extended Data Figure 1.**
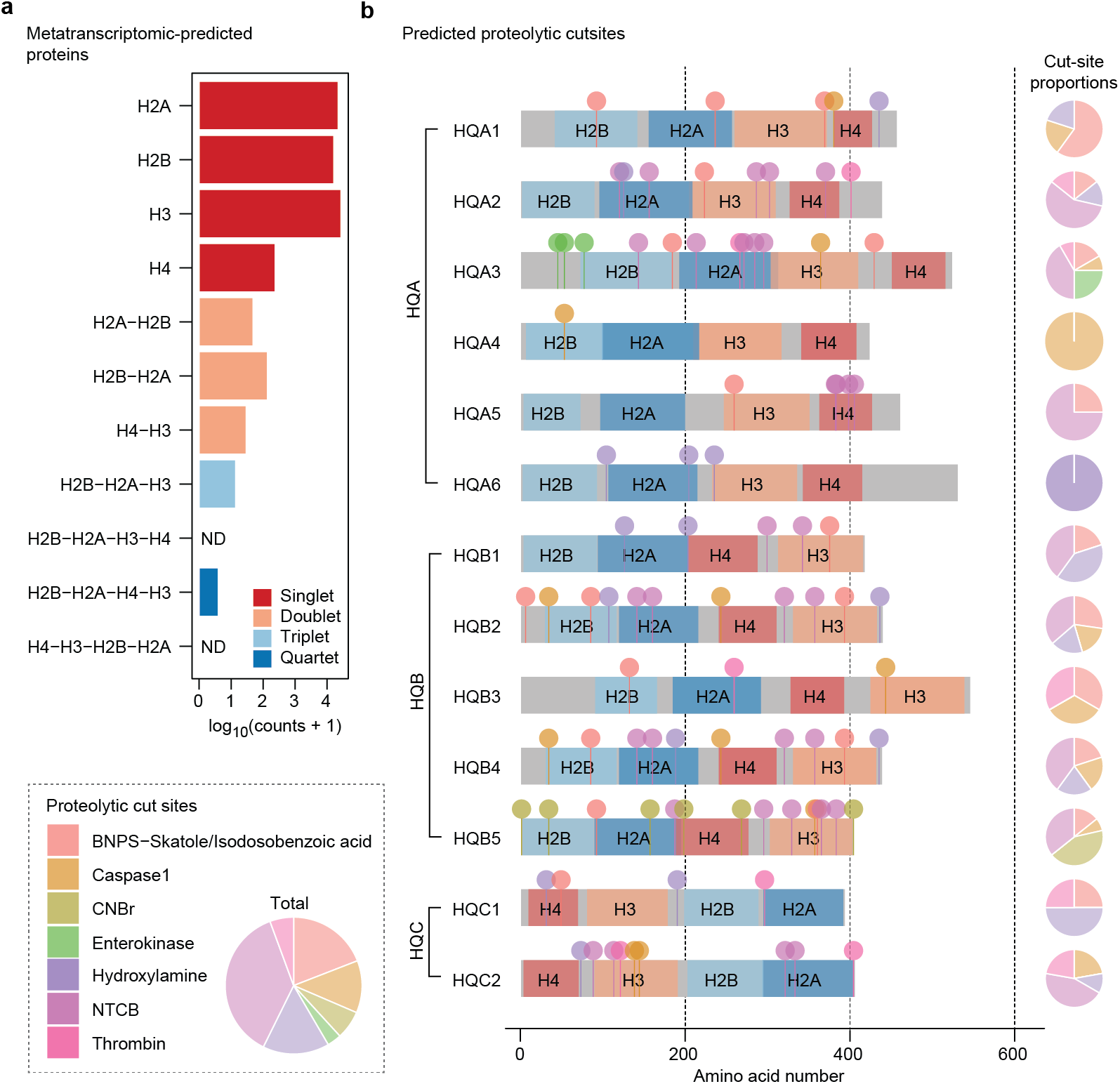
Transcription and predicted processing of viral histones indicates expression of complete repeat forms. **a**, Detection of histone repeats in metatranscriptomic data generated during the Tara Oceans Project. ND; not detected. Note that counts of individual histones will include eukaryote-derived transcripts. **b**, Predicted proteolytic cut-sites and diagrams of viral histone quadruplets. Cut-sites and cutter proportions are denoted and were predicted using ExPASy PeptideCutter.

**Extended Data Figure 2.**
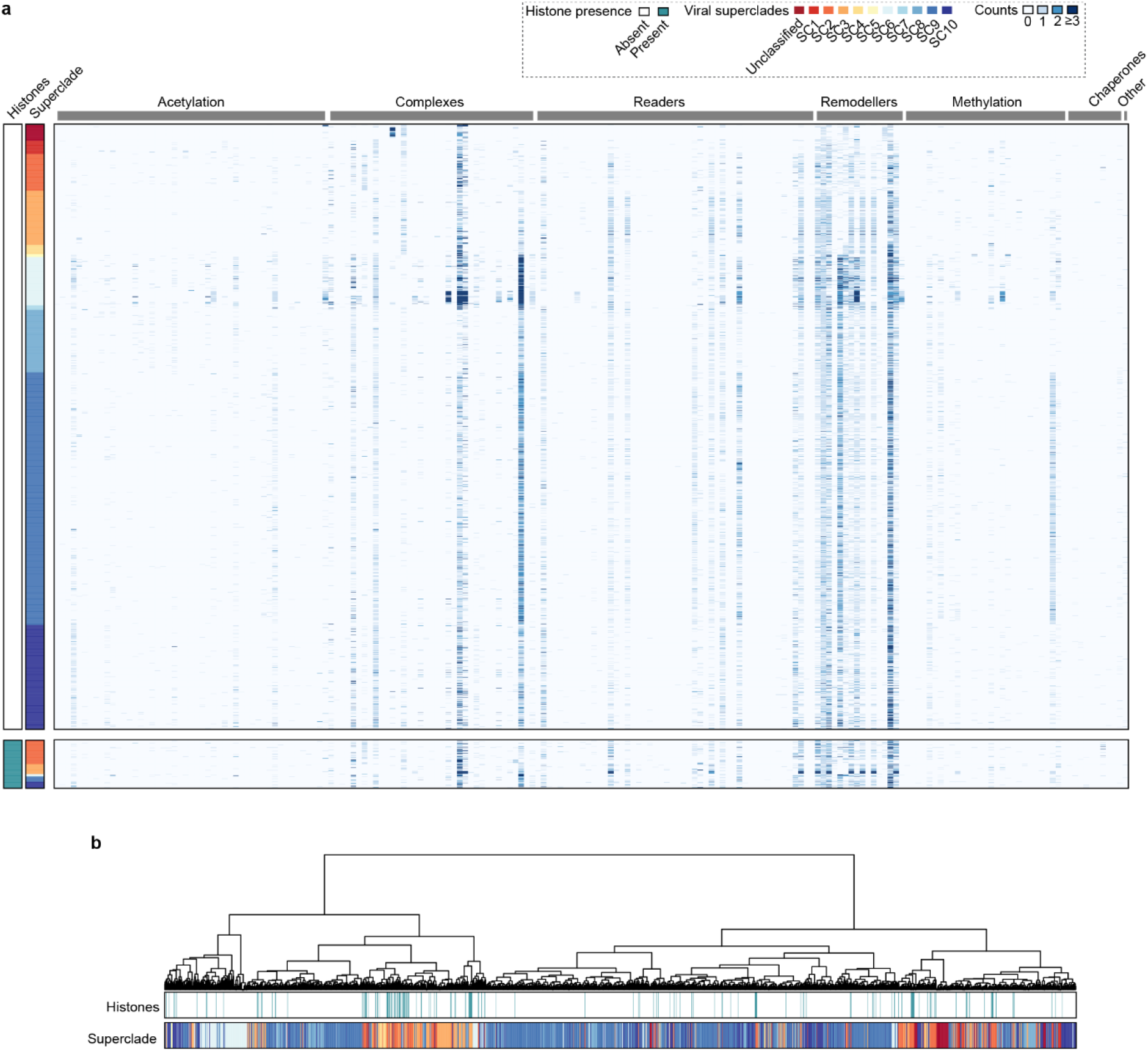
Chromatin machinery does not show a co-association with histones in Nucleocytoviricota genomes. **a**, A heatmap showing the prevalence of chromatin-associated proteins across viral genomes and metagenomes based on proteins curated by Grau-Bové et al.^44^ Histone presence and viral superclade classifications are denoted coloured strips and protein annotations are available from FigShare. **b**, Hierarchical clustering of viral taxa based on chromatin machinery repertoires. Clustering was conducted using the Ward D2 method based on correlation distances.

**Extended Data Figure 3.**
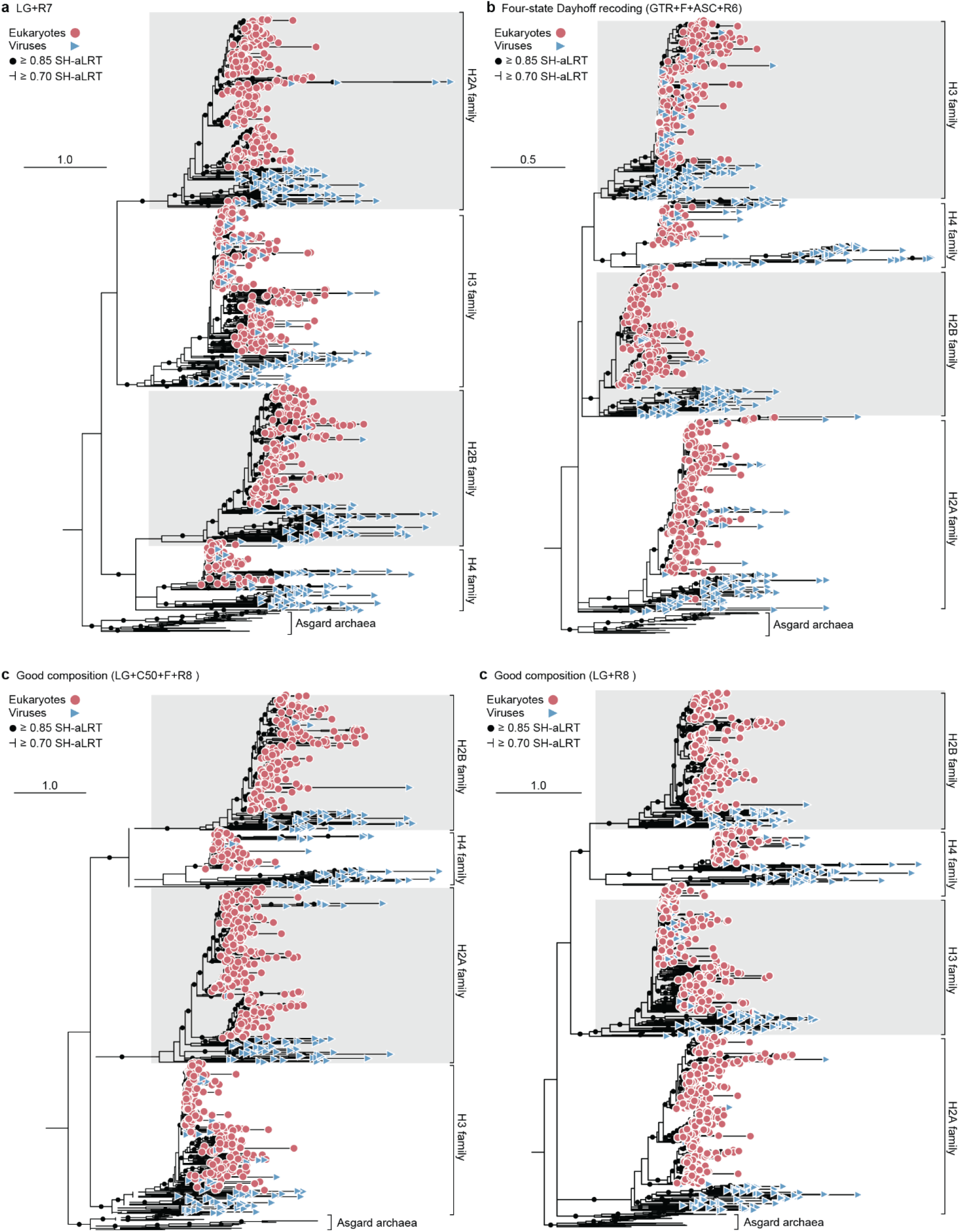
Histone phylogenies are robust to model selection but vary when accounting for compositional bias and after amino acid recoding. ML histone phylogenies generated using the same alignment as the tree in Figure 2a but using non-mixture models (**a**), after four state-Dayhoff alignment recoding (**b**), and non-mixture and mixture models after removing compositionally biased sequences (**c, d**). Statistical support was generated using SH-aLRT (*n* = 1,000) and substitution models were selected using ModelFinder and are denoted in parentheses. Full phylogenies can be viewed at https://itol.embl.de/shared/2V0SeF3AOvgMe.

**Extended Data Figure 4.**
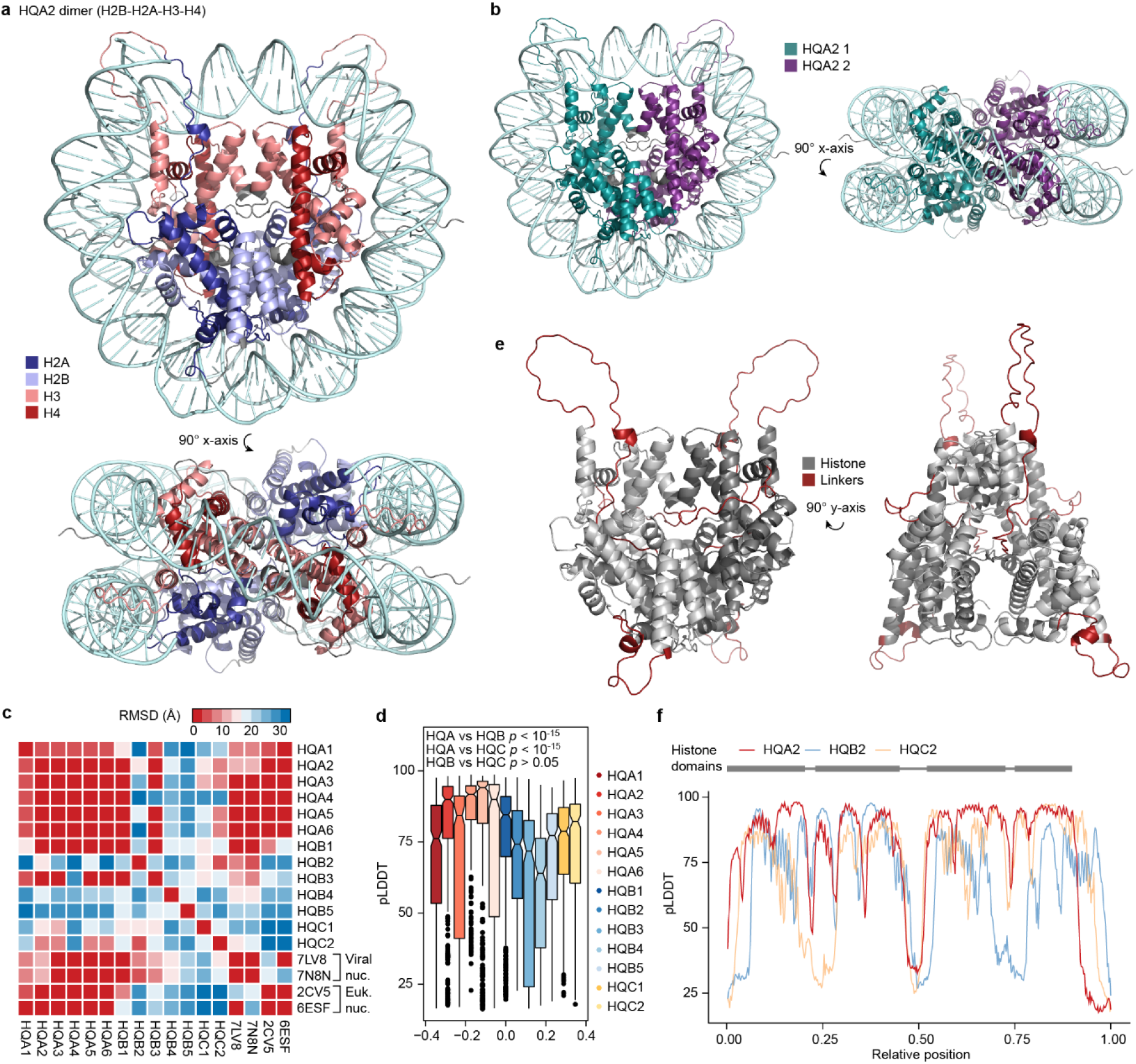
AlphaFold-predicted structures of histone quadruplet dimers reveal a capacity for nucleosome formation. The predicted structure of a histone quadruplet dimer using HQA2 as an example. Individual histone domains (**a**) or monomers (**b**) have been coloured and DNA from the structure of a *Xenopus laevis* nucleosome (PDB 6ESF) has been superimposed. **c**, Heatmap depicting root mean square deviation (RMSD) values for noted histone structures. **d**, pLDDT (predicted local distance difference test) confidence metrics for individual residues in the predicted structures of each of the histone quadruplets. Statistical comparisons of the mean confidence values were made between quadruplet types using contrasted ANOVAs after Bonferroni correction. HQA vs HQB (*p* < 2x10^-16^, F = 96.637, df = 1), HQA vs HQC (*p* < 2x10^-16^, F = 189.030, df = 1), HQB vs HQC (*p* = 0.0394, F = 4.244, df = 1). **e**, Predicted HQA2 structure with amino acid linkers highlighted in red. **f**, pLDDT confidence values across the structures of HQA2, HQB2, and HQC2. Decreases in confidence scores roughly correspond to linkers and histone domain turns. The approximate positions of the histone domains and linkers are shown. Note that the disordered C-terminus of HQA2 has been excluded from structure images.

**Extended Data Figure 5.**
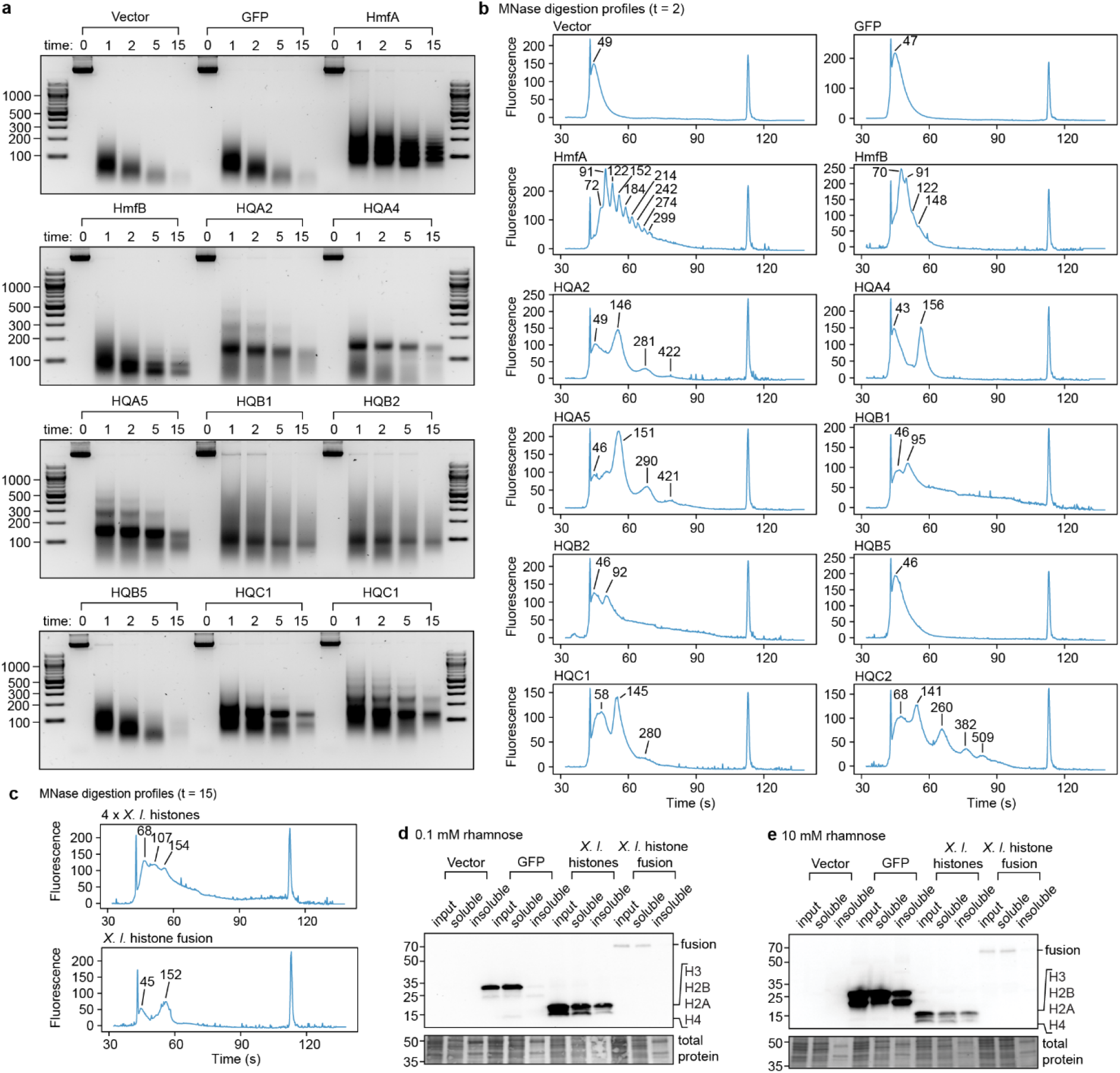
Micrococcal nuclease digestion profiles and the impact of histone fusion of protein solubility. **a**, Gel electrophoresis of genomic DNA from *E. coli* after 1, 2, 5, and 15 minutes of micrococcal nuclease (MNase) digestion during the expression of different vectors. **b**, BioAnalyzer size spectra for genomic DNA after two minutes of MNase digestion (see Fig. 2b). Peaks and their associated fragment sizes in base pairs are noted. **c**, BioAnalyzer size spectra for genomic DNA after 15 minutes of MNase digestion in the presence of individual or fused *X. laevis* histones. **d-e**, Anti-6xHis immunoblots of input, soluble, and insoluble protein fractions from *E. coli* expressing GFP or *X. laevis* histones induced by 0.1 mM (**d**) and 10 mM (**e**) rhamnose. Each experiment was repeated at least twice independently with equivalent results.

**Extended Data Figure 6.**
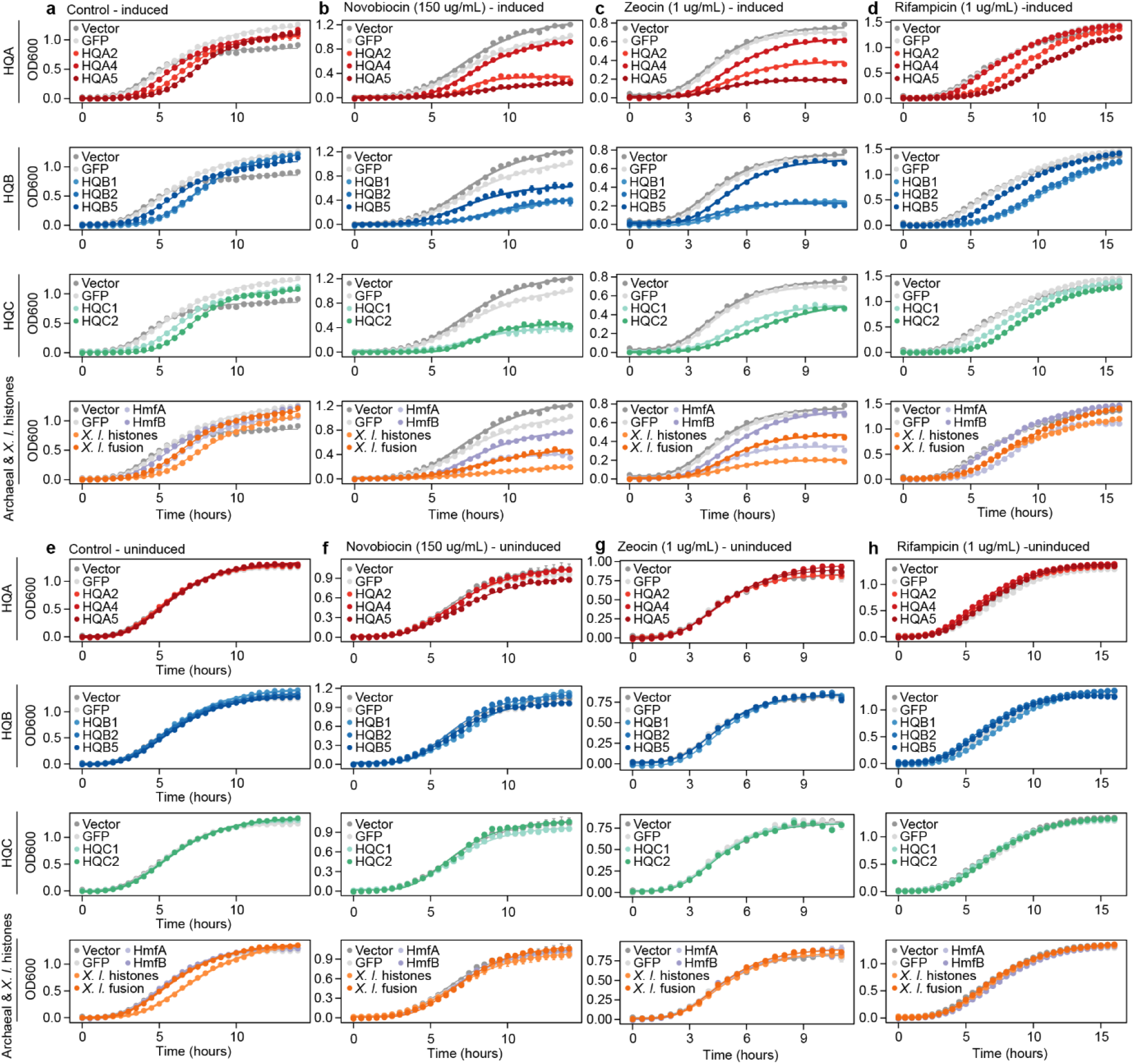
Complete *Escherichia coli* growth curves from which growth parameters were inferred. OD_600_-based growth curves for *E. coli* expressing different constructs when induced (**a-d**) or uninduced (**e-h**) under normal conditions (**a, e**) or in the presence of novobiocin (**b, f**), zeocin (**c, g**), or rifampicin (**d, h**). Data were fit to sigmoidal models. Points represent values averaged across three replicates and error bars represent the standard error of the mean. Note that the error bars are often very small and can be difficult to visualize. Each experiment was repeated at least three times independently with equivalent results.

**Table S1.** Constrained histone phylogenies and topology testing.

| Constraint<sup>1</sup> | -Log likelihood | $\Delta$ Log likelihood<sup>2</sup> | *p* value (AU test) |
| --- | --- | --- | --- |
| H2A-H2B | -111503.4297 | 108.66 | 0.249 |
| H2A-H2B-H3 | -111473.6117 | 78.838 | 0.375 |
| H2A-H2B-H4 | -111528.3568 | 133.58 | 0.187 |
| H2A-H3-H4 | -111448.9437 | 54.17 | 0.455 |
| H2A-H4 | -111506.1031 | 111.33 | 0.257 |
| H2B-H3 | -111452.709 | 57.935 | 0.467 |
| H2B-H3-H4 | -111438.9424 | 44.168 | 0.483 |
| H3-H4 | -111484.6786 | 89.905 | 0.312 |
| H2A – all repeats | -45863.28644 | 11.774 | 0.448 |
| H2A – singlets | -45983.15426 | 131.64 | 0.00852 |
| H2A – doublets | -46111.718 | 260.21 | 3.73e-05 |
| H2A – triplets | -45854.68443 | 3.1725 | 0.538 |
| H2A – quadruplets | -45917.38789 | 65.876 | 0.127 |
| H2B – all repeats | -38498.74644 | 5.1021 | 0.511 |
| H2B – singlets | -38496.98507 | 3.3408 | 0.563 |
| H2B – doublets | -38530.68588 | 37.042 | 0.225 |
| H2B – triplets | -38504.12216 | 10.478 | 0.457 |
| H2B – quadruplets | -38548.68764 | 55.043 | 0.117 |
| H3 – all repeats | -51377.08679 | 8.3113 | 0.475 |
| H3 – singlets | -51378.29098 | 9.5155 | 0.457 |
| H3 – doublets | -51373.4709 | 4.6955 | 0.493 |
| H3 – triplets | -51383.42459 | 14.649 | 0.422 |
| H3 – quadruplets | -51453.40845 | 84.633 | 0.0852 |
| H4 – all repeats | -17407.47899 | 5.8677 | 0.482 |
| H4 – singlets | -17462.15932 | 60.548 | 0.019 |
| H4 – doublets | -17436.31786 | 34.707 | 0.139 |
| H4 – quadruplets | -17488.58268 | 86.971 | 0.000159 |

<sup>1</sup> Tree topologies can be viewed at https://itol.embl.de/shared/wcmaHN1Y8dVO.
<sup>2</sup> Change in likelihood relative to the best unconstrained tree.

## References

1. Cavalier-Smith, T. Origin of the cell nucleus, mitosis and sex: roles of intracellular coevolution. Biol Direct 5, 7 (2010).

2. Luger, K., Dechassa, M. L. & Tremethick, D. J. New insights into nucleosome and chromatin structure: an ordered state or a disordered affair? Nat Rev Mol Cell Biol 13, 436–447 (2012).

3. Sandman, K. & Reeve, J. N. Archaeal histones and the origin of the histone fold. Current Opinion in Microbiology 9, 520–525 (2006).

4. Malik, H. S. & Henikoff, S. Phylogenomics of the nucleosome. Nat Struct Mol Biol 10, 882–891 (2003).

5. Brunk, C. F. & Martin, W. F. Archaeal histone contributions to the origin of eukaryotes. Trends in Microbiology 27, 703–714 (2019).

6. Talbert, P. B., Armache, K.-J. & Henikoff, S. Viral histones: pickpocket’s prize or primordial progenitor? Epigenetics & Chromatin 15, 21 (2022).

7. Erives, A. J. Phylogenetic analysis of the core histone doublet and DNA topo II genes of Marseilleviridae: evidence of proto-eukaryotic provenance. Epigenetics & Chromatin 10, 55 (2017).

8. Thomas, V. et al. Lausannevirus, a giant amoebal virus encoding histone doublets: a giant virus encoding histone doublets. Environmental Microbiology 13, 1454–1466 (2011).

9. Liu, Y. et al. Virus-encoded histone doublets are essential and form nucleosome-like structures. Cell 184, 4237–4250.e19 (2021).

10. Valencia-Sánchez, M. I. et al. The structure of a virus-encoded nucleosome. Nat Struct Mol Biol 28, 413–417 (2021).

11. Yoshikawa, G. et al. Medusavirus, a novel large DNA virus discovered from hot spring water. J Virol 93, e02130–18 (2019).

12. Roger, A. J., Susko, E. & Leger, M. M. Evolution: reconstructing the timeline of eukaryogenesis. Current Biology 31, R193–R196 (2021).

13. Schulz, F. et al. Giant virus diversity and host interactions through global metagenomics. Nature 578, 432–436 (2020).

14. Schulz, F., Abergel, C. & Woyke, T. Giant virus biology and diversity in the era of genome-resolved metagenomics. Nat Rev Microbiol 20, 721–736 (2022).

15. Irwin, N. A. T., Pittis, A. A., Richards, T. A. & Keeling, P. J. Systematic evaluation of horizontal gene transfer between eukaryotes and viruses. Nat Microbiol 7, 327–336 (2021).

16. Guglielmini, J., Woo, A. C., Krupovic, M., Forterre, P. & Gaia, M. Diversification of giant and large eukaryotic dsDNA viruses predated the origin of modern eukaryotes. Proc. Natl. Acad. Sci. U.S.A. 116, 19585–19592 (2019).

17. Bryson, T. D. et al. A giant virus genome is densely packaged by stable nucleosomes within virions. Molecular Cell 82, 4458–4470.e5 (2022).

18. Carradec, Q. et al. A global ocean atlas of eukaryotic genes. Nat Commun 9, 373 (2018).

19. Penev, P. I., Alvarez-Carreño, C., Smith, E., Petrov, A. S. & Williams, L. D. TwinCons: Conservation score for uncovering deep sequence similarity and divergence. PLoS Comput Biol 17, e1009541 (2021).

20. Ashkenazy, H. et al. ConSurf 2016: an improved methodology to estimate and visualize evolutionary conservation in macromolecules. Nucleic Acids Res 44, W344–W350 (2016).

21. Bannister, A. J. & Kouzarides, T. Regulation of chromatin by histone modifications. Cell Res 21, 381–395 (2011).

22. Jumper, J. et al. Highly accurate protein structure prediction with AlphaFold. Nature 596, 583–589 (2021).

23. Tsompana, M. & Buck, M. J. Chromatin accessibility: a window into the genome. Epigenetics & Chromatin 7, 33 (2014).

24. Rojec, M., Hocher, A., Stevens, K. M., Merkenschlager, M. & Warnecke, T. Chromatinization of Escherichia coli with archaeal histones. eLife 8, e49038 (2019).

25. Mattiroli, F. et al. Structure of histone-based chromatin in Archaea. Science 357, 609–612 (2017).

26. Luger, K., Mäder Armin W., Richmond Robin K., Sargent David F., & Richmond Timothy J. Crystal structure of the nucleosome core particle at 2.8 Å resolution. Nature 389, 251–260 (1997).

27. Burgess, R. J. & Zhang, Z. Histone chaperones in nucleosome assembly and human disease. Nat Struct Mol Biol 20, 14–22 (2013).

28. Shim, Y., Duan, M.-R., Chen, X., Smerdon, M. J. & Min, J.-H. Polycistronic coexpression and nondenaturing purification of histone octamers. Analytical Biochemistry 427, 190–192 (2012).

29. Thomas, J. G. & Baneyx, F. Protein misfolding and inclusion body formation in recombinant Escherichia coli cells overexpressing heat-shock proteins. Journal of Biological Chemistry 271, 11141–11147 (1996).

30. Clark, D. J. & Felsenfeld, G. Formation of nucleosomes on positively supercoiled DNA. The EMBO Journal 10, 387–395 (1991).

31. Stevens, K. M. et al. Histone variants in Archaea and the evolution of combinatorial chromatin complexity. Proc. Natl. Acad. Sci. U.S.A. 117, 33384–33395 (2020).

32. Eme, L. et al. Inference and reconstruction of the heimdallarchaeial ancestry of eukaryotes. Nature (2023) doi:10.1038/s41586-023-06186-2.

33. The UniProt Consortium. UniProt: a worldwide hub of protein knowledge. Nucleic Acids Research 47, D506–D515 (2019).

34. Simão, F. A., Waterhouse, R. M., Ioannidis, P., Kriventseva, E. V. & Zdobnov, E. M. BUSCO: assessing genome assembly and annotation completeness with single-copy orthologues. Bioinformatics 31, 3210–3212 (2015).

35. Fu, L., Niu, B., Zhu, Z., Wu, S. & Li, W. CD-HIT: accelerated for clustering the next-generation sequencing data. Bioinformatics 28, 3150–3152 (2012).

36. Buchfink, B., Reuter, K. & Drost, H.-G. Sensitive protein alignments at tree-of-life scale using DIAMOND. Nat Methods 18, 366–368 (2021).

37. Altschul, S. F., Gish, W., Miller, W., Myers, E. W. & Lipman, D. J. Basic Local Alignment Search Tool. Journal of Molecular Biology 215, 403–410 (1990).

38. Katoh, K. & Standley, D. M. MAFFT Multiple Sequence Alignment Software Version 7: improvements in performance and usability. Molecular Biology and Evolution 30, 772–780 (2013).

39. Capella-Gutiérrez, S., Silla-Martínez, J. M. & Gabaldón, T. trimAl: a tool for automated alignment trimming in large-scale phylogenetic analyses. Bioinformatics 25, 1972–1973 (2009).

40. Kalyaanamoorthy, S., Minh, B. Q., Wong, T. K. F., Von Haeseler, A. & Jermiin, L. S. ModelFinder: fast model selection for accurate phylogenetic estimates. Nat Methods 14, 587–589 (2017).

41. Boeckmann, B. The SWISS-PROT protein knowledgebase and its supplement TrEMBL in 2003. Nucleic Acids Research 31, 365–370 (2003).

42. Mistry, J., Finn, R. D., Eddy, S. R., Bateman, A. & Punta, M. Challenges in homology search: HMMER3 and convergent evolution of coiled-coil regions. Nucleic Acids Research 41, e121–e121 (2013).

43. Hyatt, D. et al. Prodigal: prokaryotic gene recognition and translation initiation site identification. BMC Bioinformatics 11, 119 (2010).

44. Grau-Bové, X. et al. A phylogenetic and proteomic reconstruction of eukaryotic chromatin evolution. Nat Ecol Evol 6, 1007–1023 (2022).

45. Suzuki, R. & Shimodaira, H. Pvclust: an R package for assessing the uncertainty in hierarchical clustering. Bioinformatics 22, 1540–1542 (2006).

46. Guindon, S. et al. New algorithms and methods to estimate maximum-likelihood phylogenies: assessing the performance of PhyML 3.0. Systematic Biology 59, 307–321 (2010).

47. Nguyen, L.-T., Schmidt, H. A., Von Haeseler, A. & Minh, B. Q. IQ-TREE: a fast and effective stochastic algorithm for estimating maximum-likelihood phylogenies. Molecular Biology and Evolution 32, 268–274 (2015).

48. Shimodaira, H. An approximately unbiased test of phylogenetic tree selection. Systematic Biology 51, 492–508 (2002).

49. Letunic, I. & Bork, P. Interactive Tree Of Life (iTOL) v5: an online tool for phylogenetic tree display and annotation. Nucleic Acids Research 49, W293–W296 (2021).

50. Mirdita, M. et al. ColabFold: making protein folding accessible to all. Nat Methods 19, 679–682 (2022).

51. Gasteiger, E. ExPASy: the proteomics server for in-depth protein knowledge and analysis. Nucleic Acids Research 31, 3784–3788 (2003).

52. Steinegger, M. & Söding, J. MMseqs2 enables sensitive protein sequence searching for the analysis of massive data sets. Nat Biotechnol 35, 1026–1028 (2017).

